# Tirzepatide restricts obesity-related tumor growth by reversing metabolic dysregulation and rescuing CD8+ T cell function

**DOI:** 10.1101/2024.01.20.576484

**Authors:** Elaine M. Glenny, Michael F. Coleman, Genevieve T. Clutton, Brian P. Riesenberg, Alyssa N. Ho, Tori L. McFarlane, Violet A. Kiesel, Amanda L. Kucinskas, Emma G. Bailey, Katherine E. Sanchez, Jibin Zeng, Ye Wang, Hannah M. Malian, Claire E. Gates, Fangxin Chen, Ruihan Xu, C. Alex Holt, Evan M. Paules, Erin D. Giles, Jessica E. Thaxton, Michael Pollak, Stephen D. Hursting

**Author notes:** Co-corresponding authors Stephen D. Hursting, PhD, MPH Michael Pollak, MD Jessica E. Thaxton, PhD, MsCR.

## Abstract

Obesity, an established risk and progression factor for at least 13 cancer types, is highly prevalent globally, and effective strategies to mitigate the burden of obesity-related cancer are urgently needed. We investigated whether tirzepatide, a widely used incretin-mimetic drug that induces substantial weight loss, offers anticancer benefits. Across 3 tumor models, we demonstrate that chronic tirzepatide treatment reverses diet-induced increases in body weight and fat mass, systemic metabolic perturbations, and tumor growth. We also showed that the anticancer activity of tirzepatide does not involve direct effects on the neoplastic cells used, which lack incretin receptor expression. The anticancer actions of tirzepatide require the reversal of both the metabolic dysregulation and hyporesponsiveness of CD8+ tumor infiltrating lymphocytes evident in obesity. Our findings establish tirzepatide as a promising compound for intercepting obesity-related cancers.

## Introduction

The global epidemic of obesity^1^, which is a leading risk and progression factor for many common cancers^2^, motivates the development of effective strategies to mitigate this dual public health crisis. Obesity drives multiple metabolic alterations, including hyperglycemia, hyperleptinemia, hyperinsulinemia, as well as low-grade chronic inflammation^3^. These perturbations foster an environment in which cancer cells receive mitogenic signals that not only accelerate proliferation, but also lead to increased uptake and utilization of macronutrients to the detriment of competing intratumoral immune cells^4–6^. Obesity further impairs antitumor immunity by engendering an immunosuppressive environment that protects tumor cells from the cytotoxic effects of CD8+ tumor infiltrating lymphocytes (TILs)^6–9^.

Bariatric surgery is currently the most effective long-term weight loss strategy in patients with severe obesity, achieving >25% average total weight loss at 5 years accompanied by metabolic and immune reprogramming and a sustained 30-50% risk reduction in cancers associated with obesity^10,11^. However, bariatric surgery is expensive and only 1% of qualifying individuals in the United States undergo the procedure^12^. Lifestyle modifications through diet and/or exercise are generally accessible but typically result in only modest short-term weight loss that is challenging for most people to sustain^13,14^. Incretin-based pharmacotherapies, including semaglutide a glucagon-like peptide (GLP)-1 receptor agonist (GLP1RA) and tirzepatide a dual GLP-1 and glucose-dependent insulinotropic polypeptide (GIP) receptor agonist, are increasingly prescribed for the management of obesity and/or type 2 diabetes^15^. These drugs cause weight loss largely by acting in the central nervous system to suppress appetite and reduce food intake^16^. Tirzepatide, a highly effective weight loss drug approved by the US Food and Drug Administration in 2022, achieves an average sustained weight loss of 22.5% among adults with obesity^17,18^, reduces the risk of several chronic diseases, including type 2 diabetes^19^ and cardiovascular disease^20^, and is safe and well tolerated over >3 years^21^. Incretin-mimetic agents offer a promising, yet understudied, interception strategy for obesity-related cancers.

A study of patients with type 2 diabetes followed for 15 years showed that GLP1RA use, relative to insulin use, was associated with lower risk of 10 of 13 obesity-related cancers, particularly endometrial cancer in women and colon cancer in men^22^. In other studies of patient cohorts with type 2 diabetes, associations of GLP1RA use and cancer risk were mixed^23–28^. These retrospective pharmacoepidemiologic studies must be interpreted with caution due to lack of randomization, particularly in the absence of supporting laboratory data and understanding of underlying mechanisms. Early laboratory studies of first-generation GLP1RAs and cancer progression showed inconsistent results^29^, while studies of tirzepatide are sparse and have not elucidated the underlying biology^30,31^. In experimental systems using next-generation incretin mimetics, the extent of protection against obesity-driven tumor growth follows the extent of weight loss achieved^32^. Herein, using multiple obesity-responsive syngeneic mouse transplant models of cancer, we interrogated the effects of tirzepatide on systemic metabolism, antitumor immunity, and tumor progression.

## Results

### Tirzepatide-associated weight loss in obese mice limits tumor growth and reverses metabolic perturbations

We first determined if chronic administration of tirzepatide reduces obesity-driven tumor growth in the immunogenic MC38 cancer model^6^. Diet-induced obese (DIO) mice received subcutaneous injections of either tirzepatide (to promote weight loss) or vehicle (to maintain obesity) for 3 weeks and were then transplanted with MC38 cancer cells (**Figure 1A**). Control diet-fed mice were injected with vehicle and served as never-obese controls. A dose-finding study in DIO mice was initially conducted to establish a tirzepatide regimen that achieves the ∼22% weight loss observed in human studies^17,18,21^. Weight loss was assessed over a range of doses (1-30 nmol/kg) at different frequencies of administration (daily, every other day [q.o.d.], and three times per week) for a 4-week duration (**Supplementary Figure 1**). Based on these results, tirzepatide was administered in the current study q.o.d. using a progressive dose escalation strategy (3-10 nmol/kg), with ∼25% weight loss achieved by day 21 and maintained (**Figure 1B**). As previously reported, the weight loss coincided with decreased caloric intake in the setting of unrestricted food availability, with the largest reductions in food intake occurring during the initial 4 weeks of tirzepatide treatment (**Figure 1B-C**)^33,34^.

**Figure 1.**
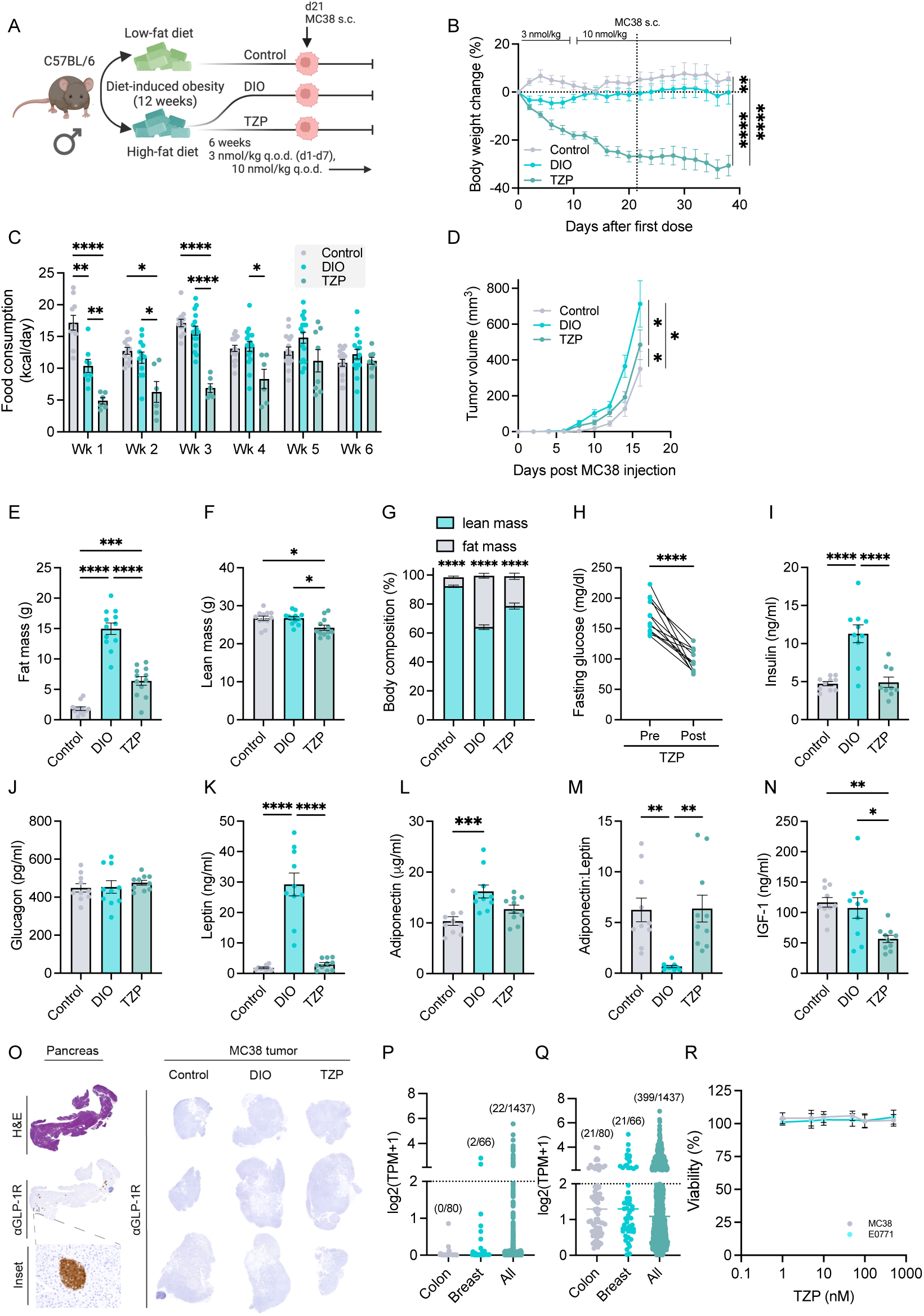
Tirzepatide (TZP) limits tumor growth in obesity. **(A)** Study design for chronic TZP treatment in mice bearing subcutaneous MC38 tumors. **(B)** Percent change in body weight over 6 weeks with q.o.d. subcutaneous TZP or vehicle injections (n=12/12/12). **(C)** Average calories consumed per mouse for two consecutive 24-hour intervals each week (n=16/16/8). **(D)** MC38 tumor volume over time (n=12/12/12). **(E)** Fat mass (n=12/12/12), **(F)** lean mass (n=12/12/12), and **(G)** body composition (n=12/12/12) after 20 days of treatment with TZP or vehicle. **(H)** Fasting blood glucose prior to and following 20 days of TZP (n=12/12/12). Fasting serum concentrations of **(I)** insulin (n=10/10/9), **(J)** glucagon (n=10/10/10), **(K)** leptin (n=9/10/10), **(L)** adiponectin (n=10/10/10), **(M)** adiponectin:leptin (n=10/10/10), and **(N)** IGF-1 (n=10/10/10). **(O)** Representative immunohistochemistry staining for Glp1r on pancreas and MC38 tumor sections. Expression levels of *GLP1R* **(P)** and *GIPR* **(Q)** in CCLE RNAseq data. **(R)** MC38 and E0771 cell viability assessed by MTT assay (n=5/group) after 72 h TZP treatment at indicated doses. Data are represented as mean ± SEM. Statistical significance determined by repeated measures two-way ANOVA with Tukey’s multiple comparisons test (B, G), one-way ANOVA with Tukey’s multiple comparisons test (C, E-F, I-N), linear regression with FDRq correction (D), paired two-sided Student’s t-test (H), or repeated measures one-way ANOVA with Dunnett’s multiple comparisons test (R).

Along with inducing weight loss, chronic tirzepatide administration to DIO mice significantly restricted tumor growth (**Figure 1D**) coincident with an improved metabolic profile prior to cancer cell allograft. Specifically, tirzepatide drove substantial loss of total body mass, with 78% of the weight loss attributed to fat mass, and reduction in fasting blood glucose levels (**Figure 1E-H**). Several DIO-induced systemic metabolic perturbations were also reversed by tirzepatide to resemble phenotypically never-obese controls, particularly i) leptin, which decreased >9-fold; ii) insulin, which decreased 2-fold; and iii) adiponectin/leptin ratio, which increased >9-fold (**Figure 1I-N**). Thus, tirzepatide reduces obesity and markedly improves the metabolic milieu of DIO mice in association with reversal of obesity-driven MC38 tumor growth.

To establish generalizability across sex and tumor type, we evaluated additional models. Obese female mice were treated with similarly escalating doses of tirzepatide (3-30 nmol/kg) or 30% chronic calorie restriction, a gold-standard murine intervention to limit metabolic disease and tumor growth. Tirzepatide and CCR both corrected metabolic abnormalities of obese mice, reducing body weight and fat mass (**Supplementary Figure 2A-C**) and normalized the levels of metabolic hormones and glucose in circulation (**Table S1**), with more substantial metabolic effects elicited by CCR than by tirzepatide. Obesity-driven growth of E0771 tumors was blunted by both tirzepatide and CCR (**Supplementary Figure 2D**). Similarly, tirzepatide treatment, relative to vehicle treatment, in obese mice, resulted in a 40% reduction in transplanted ER+ Py230 tumor volume (**Supplementary Figure 2E**). Finally, terminal body weight was significantly associated with tumor burden in both breast cancer models (**Supplementary Figure 2F-G**). Thus, the mitigation of obesity-driven tumor growth was consistent across multiple tumor models and both sexes.

### Tirzepatide efficacy is independent of direct GLP1R or GIPR engagement on MC38 tumor cells

To address the mechanism underlying the antitumor activity of tirzepatide, we investigated the possibility that the drug directly engages with GLP1R or GIPR on MC38 cancer cells. Immunohistochemical analysis revealed that GLP1R protein (**Figure 1O**) was not expressed in MC38 tumors (no validated antibody currently exists for murine GIPR^35^), consistent with our analysis of the Cancer Cell Line Encyclopedia database^36^ showing 0/80 human colorectal cancer cell lines and 2/66 human breast cancer cell lines (and only 22/1437 of all human cancer cell lines) express *GLP1R* at appreciable levels^37^ (**Figure 1P**). While 21/80 (25%) human colorectal cancer cell lines and 21/66 human breast cancer cell lines (and 399/1437 of total cancer cell lines) express low but discernible *GIPR* (**Figure 1Q**), this is likely trivial in murine models where GIPR is only engaged at high concentrations of tirzepatide (>30 nmol/kg)^38^. In addition, MC38 and E0771 cell viability in vitro was unaffected by exposure to tirzepatide across a 1–500 nM dose range (**Figure 1R**). Accordingly, we dismissed direct effects of tirzepatide on cancer cells as an explanation for the observed inhibition of allograft growth.

### CD8+ TILs restrain tumor growth in tirzepatide-treated DIO mice

We used bulk RNAseq of MC38, E0771, and Py230 tumors to reveal pathways and processes altered by obesity (relative to control) that were normalized with chronic administration of tirzepatide. Notably, features of antitumor immunity were consistently suppressed by obesity but rescued with tirzepatide (**Supplementary Figure 2H**).

Obesity promotes an immunosuppressive tumor microenvironment that profoundly impairs the ability of CD8+ TILs to effectively respond to and kill cancer cells^6–8^. We hypothesized that tirzepatide-induced weight loss and reversal of obesity-associated metabolic derangements would ameliorate the obesity-driven CD8+ TIL dysfunction that leads to a deficit in antitumor immunity and increased tumor growth. To further investigate this hypothesis, we relied on the MC38 tumor model as it is well characterized as both obesity-responsive and immunogenic. Mice that received 21 days of vehicle or tirzepatide q.o.d., following 12 weeks of control or DIO diet, were challenged with MC38 cancer cells, continued their assigned diet and drug intervention, and were then euthanized after 14 days of tumor growth. We employed high-dimensional flow cytometry analysis of tumoral and splenic CD8+ T cells to delineate obesity-driven defects in antitumor immunity potentially rescued by tirzepatide. Uniform manifold approximation and projection (UMAP) analysis of CD8+ TILs from tumors of vehicle-treated control (never-obese) versus vehicle-treated DIO (always obese) mice identified 17 distinct clusters (**Figure 2A-B**). CD8+ TILs from vehicle-treated never-obese mice were enriched in clusters 1, 4, and 5 and were characterized by higher staining intensities for markers of activation (CD44), effector function (GZMB), and exhaustion (CD39, PD-1, TIM-3, CTLA-4) (**Figure 2C-D**). Conversely, CD8+ TILs from vehicle-treated DIO mice were enriched in clusters 15 and 17, which had considerably lower staining intensity for markers of activation, effector function, and exhaustion, and higher staining intensity for markers of non-activated and stem-like CD8+ TIL states (TCF-1, CD62L, BCL-2) (**Figure 2C-D**).

**Figure 2.**
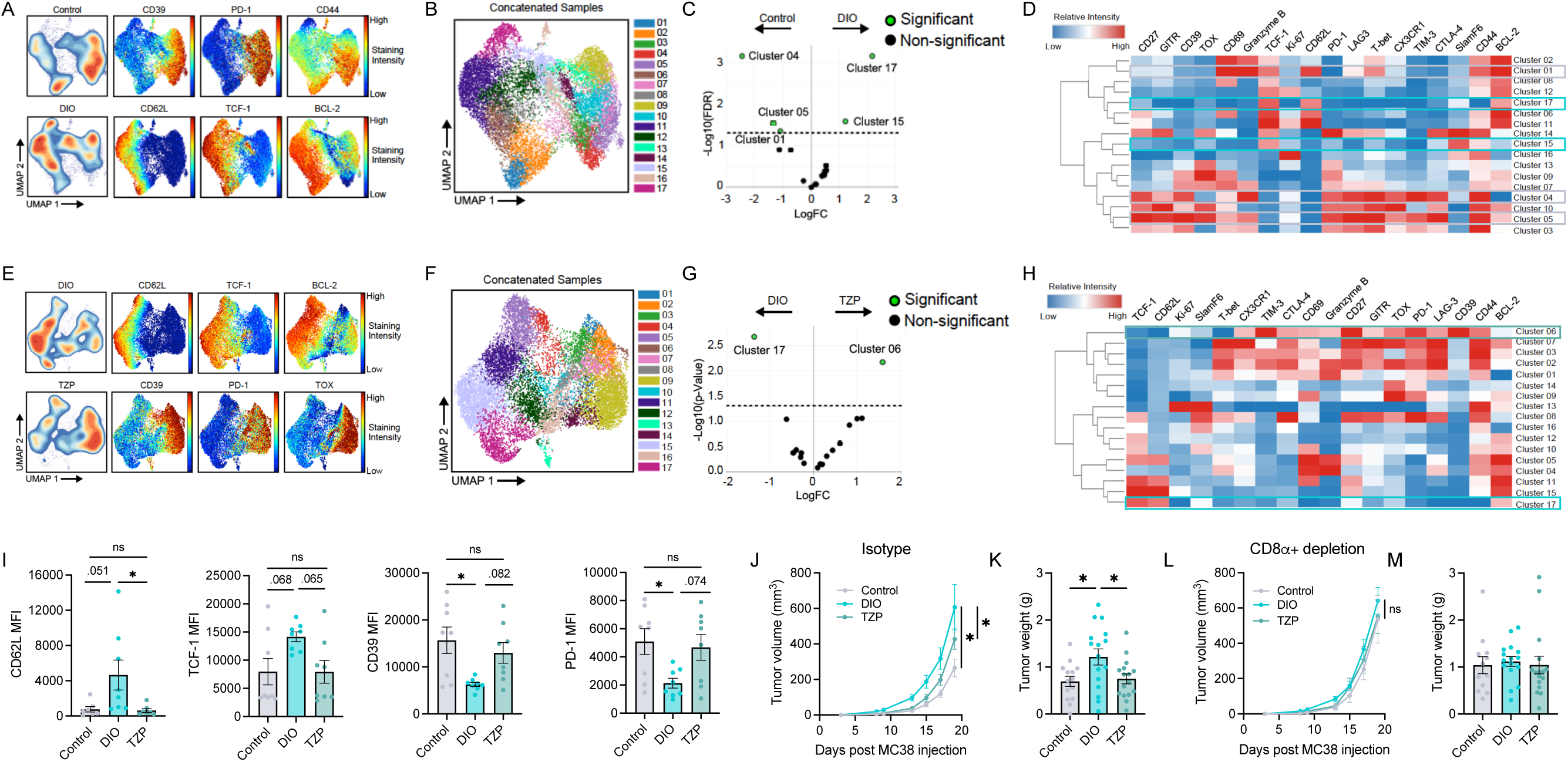
Tirzepatide (TZP) restores CD8+ T cell function to limit tumor growth in formerly obese mice. **(A)** UMAPs of high-dimensional flow cytometry on CD8+ TILs isolated from MC38 tumors of control and DIO mice at 14 days post-transplantation (n=8/8). Staining intensities of selected phenotypic markers driving UMAP projection shown. **(B)** Semi-supervised UMAP clustering of high-dimensional flow cytometry data. **(C)** Differential abundance of TILs from control and DIO mice belonging to identified clusters. **(D)** Relative staining intensities for phenotypic markers across the clusters. **(E)** UMAPs of high-dimensional flow cytometry on CD8+ TILs isolated from MC38 tumors of vehicle- or TZP-treated DIO mice at 14 days post-transplantation (n=8/8). Staining intensities of selected phenotypic markers driving UMAP projection shown. **(F)** Semi-supervised UMAP clustering of high-dimensional flow cytometry data from panel E. **(G)** Differential abundance of TILs from vehicle and TZP-treated DIO mice belonging to identified clusters. **(H)** Relative staining intensities for phenotypic markers across the clusters from panel G. **(I)** CD62L (n=7/8/7), TCF-1 (n=8/8/8), CD39 (n=8/8/8), and PD-1 (n=8/8/8) MFIs from CD8+ TILs. **(J)** MC38 tumor volume over time and **(K)** weights of isotype-treated control, DIO, and TZP mice (n=14/17/18). **(L)** MC38 tumor volume over time and **(M)** weights of αCD8α-treated control, DIO, and TZP mice (n=13/18/17). Data are represented as mean ± SEM. Statistical significance determined by edgeR differential analysis (C, G), one-way ANOVA with Tukey’s multiple comparisons test (I-M), or linear regression with FDRq correction (J, L).

We repeated the dimension reduction and clustering analysis, this time comparing CD8+ TILs from tirzepatide-treated DIO (formerly obese) mice versus vehicle-treated DIO (always obese) mice. CD8+ TILs from vehicle-treated DIO mice were enriched in cluster 17 (non-activated and stem-like TILs) and depleted in cluster 6 (activated and exhausted TILs) relative to CD8+ TILs from tirzepatide-treated DIO mice (**Figure 2E-H**). Phenotypically, CD8+ TILs from tirzepatide-treated DIO mice more closely resembled CD8+ TILs from vehicle-treated control (never-obese) mice than from vehicle-treated DIO mice, as evidenced by median fluorescent intensities (MFIs) of specific markers including CD62L, TCF-1, CD39, and PD-1 (**Figure 2I**). The mitigating effect of tirzepatide on obesity-induced suppression of CD8+ T cells was limited to TILs in the tumor microenvironment, as the immunophenotype of splenic CD8+ T cells from vehicle-versus tirzepatide-treated DIO mice was similar (**Supplementary Figure 3**). Collectively, these data suggest obesity engenders an immunosuppressive tumor environment that enriches for a population of ineffective, non-activated CD8+ TILs, and that this immunosuppressed phenotype is partially reversed with chronic tirzepatide treatment.

To assess if CD8+ TILs are essential for the anticancer benefit imparted by tirzepatide treatment in the context of obesity, we depleted CD8α+ T cells in vehicle-treated mice on the control diet, vehicle-treated DIO mice, and tirzepatide-treated DIO mice, all bearing MC38 tumors. CD8α+ T cells were systemically depleted beginning 2 days prior to MC38 transplant (**Supplementary Figure 3B-D**). In mice treated with an isotype control IgG (non-neutralizing) antibody, tirzepatide-treated DIO mice showed reduced MC38 tumor growth relative to vehicle-treated DIO mice (**Figure 2J-K**). Depletion of systemic and tumoral CD8α+ cells did not further accelerate tumor growth in obese mice whose TILs were already functionally compromised to the extent that they were not effectively restraining neoplastic proliferation. However, CD8+ T cell ablation did accelerate tumor growth in non-obese mice on the control diet and in tirzepatide-treated DIO mice, revealing a clear impairment of antitumor immunity in response to obesity which is reversible with tirzepatide (**Figure 2L-M**).

### Direct receptor agonism on CD8+ TILs does not mediate the anticancer activity of tirzepatide

GLP1R is reportedly expressed on a subset of activated T cells in various non-neoplastic contexts, and GLP1R agonism alters T cell cytokine production^39–41^. However, as this observation has yet to be tested in a cancer model, we first queried whether *GLP1R* and/or *GIPR* were detectable in CD8+ T cells in the tumor microenvironment. Publicly available CD8+ TIL single-cell sequencing data from B16 melanoma tumors (GSE245657) revealed that every CD8+ TIL cluster, with the notable exception of the CD8+ naïve T cell population, contained *GLP1R* expressing cells (**Supplementary Figure 4A-C**)^8^. However, only 1.5% of the CD8+ TIL population from either mice on the control diet or DIO mice expressed *GLP1R* (**Supplementary Figure 4A-C**). *GIPR* was not detected in any CD8+ TIL population.

Since the single-cell RNA sequencing analysis showed that *GLP1R,* but not *GIPR*, was detectable in a subset of CD8+ TILs, we next sought to determine whether direct *GLP1R* agonism on CD8+ TILs contributes to the anticancer and immunomodulating effects of tirzepatide observed herein. Congruent with published reports^39–41^, further analysis of the single-cell data (GSE245657) revealed pathways involved in T cell activation, differentiation, chemotaxis, and interferon-mediated signaling were uniformly suppressed in activated *GLP1R*+ TILs relative to activated *GLP1R*-TILs, as determined by GSEA of gene ontology biological process (GOBP) (**Supplementary Figure 4D**). Moreover, in vitro tirzepatide treatment modestly suppressed the cytotoxic function of in vitro-activated CD8+ splenocytes from tumor-bearing DIO mice, as indicated by reduced IFNγ and TNFɑ production (**Supplementary Figure 4E-G**). Collectively, these data indicate that *GLP1R* expression on CD8+TILs cells is scant, *GIPR* is undetectable, and agonism of GLP1R in CD8+ TILs is modestly immunosuppressive. Thus, direct receptor agonism on CD8+ TILs is unlikely to mediate the antitumor effects of tirzepatide in obese mice.

### Tirzepatide fails to limit MC38 tumor growth or improve CD8+ T cell response in lean mice

Since the anticancer effects of tirzepatide require CD8+ T cells but not direct receptor engagement on either tumor or CD8+ T cells, we explored the possibility that reversing obesity-associated metabolic dysfunction is key to the inhibitory effect of tirzepatide on obesity-stimulated MC38 tumor growth. We transplanted MC38 cells into lean (never-obese) mice fed an ad libitum control diet (10 kcal% fat) and then randomized the mice to receive vehicle or tirzepatide injections q.o.d. for the next 20 days (**Figure 3A**). Tirzepatide did not affect tumor growth in these lean, metabolically healthy mice despite achieving modest reductions in food intake, body weight, blood glucose, and serum levels of insulin, IGF-1, and leptin (**Figure 3B-J**). Importantly, in these lean mice, baseline hormone levels were lower, and the reductions were smaller in magnitude, than those seen in DIO mice treated with tirzepatide (**Figure 1**). To evaluate whether tirzepatide had an effect on remodeling CD8+ TIL populations in lean mice, we performed high-dimensional flow cytometry on tumoral and splenic CD8+ T cells following 14 days of MC38 tumor growth. Unlike our observations in tirzepatide-treated DIO mice, tirzepatide treatment in lean mice skewed CD8+ TILs towards a less differentiated phenotype, as evidenced by increased CD62L and SlamF6 and reduced CD44, T-bet, TOX, and CD27 (**Figure 3K-L**). Thus, neither the tumor growth inhibitory effects nor the immune-enhancing effects of tirzepatide observed in obese mice were evident in metabolically healthy lean mice. This indicates that tirzepatide enhances anticancer immunity in obese mice indirectly, by ameliorating endocrine or metabolic abnormalities associated with obesity. This is consistent with prior evidence that obesity-related endocrine abnormalities such as hyperleptinemia compromise anticancer immune function^42,43^.

**Figure 3.**
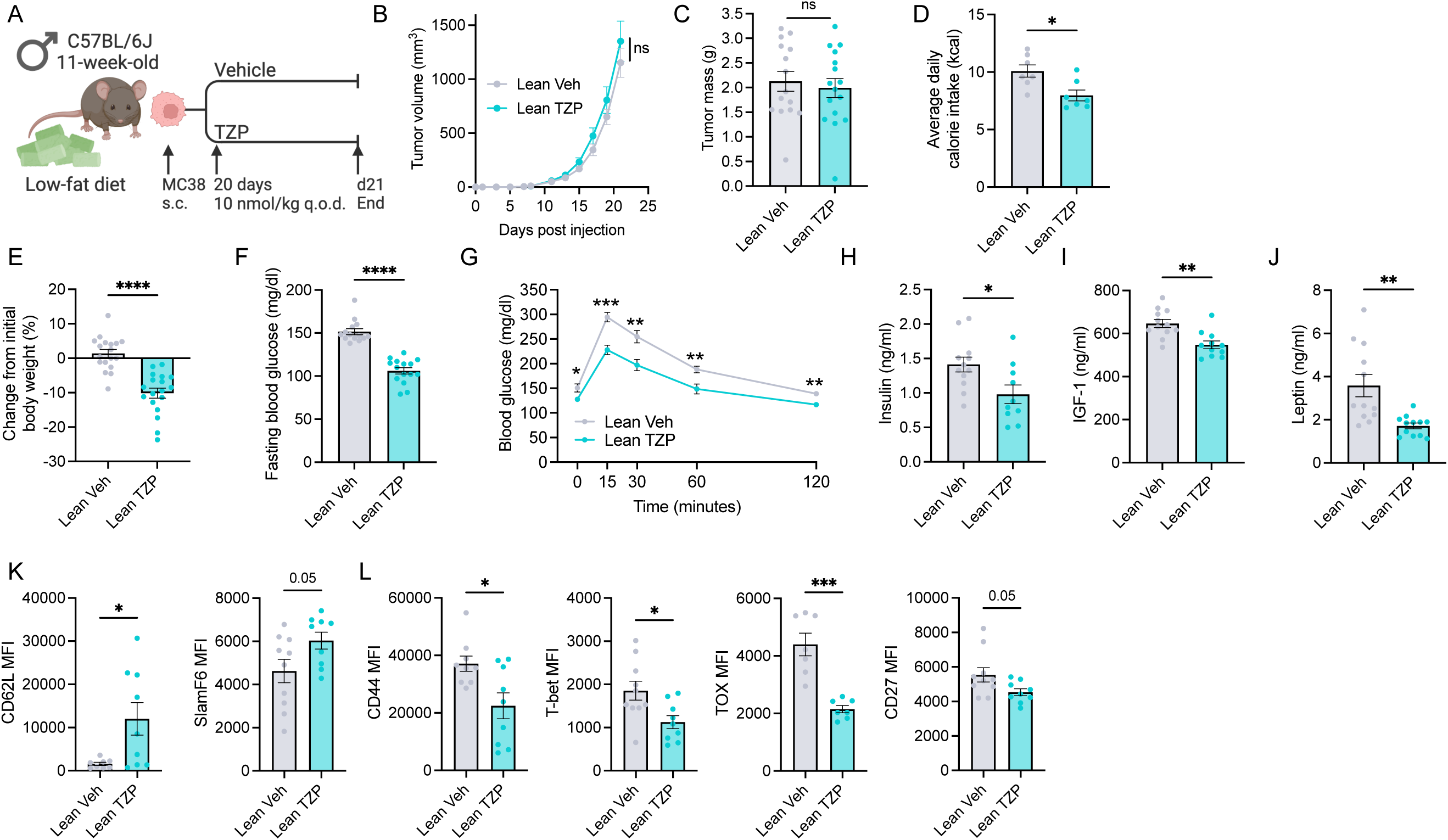
Metabolic dysfunction is essential to achieve an anticancer benefit with tirzepatide (TZP). **(A)** TZP treatment in lean mice with MC38 subcutaneous tumor transplant study design. **(B)** MC38 tumor volume over time and **(C)** tumor mass (n=15/16). **(D)** Average daily calorie intake per mouse (n=7/7). **(E)** Percent body weight change (n=15/16). **(F)** Fasting blood glucose (n=14/15). **(G)** Blood glucose concentration following intraperitoneal bolus of 1.5 mg/kg D-glucose at indicated time points (n=10/10). Fasting serum concentrations of **(H)** insulin (n=12/10), **(I)** IGF-1 (n=12/11), and **(J)** leptin (n=12/12). **(K)** CD62L (n=8/9) and SlamF6 MFIs from CD8+ TILs. **(L)** CD44 (n=9/9), T-bet (n=10/9), (TOX (n=7/7), and CD27 (n=10/9) MFIs from CD8+ TILs. Data are represented as mean ± SEM. Statistical significance determined by linear regression with FDRq correction (B) or unpaired two-sided Student’s t-test (C-L).

## Discussion

We demonstrate that chronic tirzepatide treatment reverses obesity-associated increases in body weight, fat mass, and systemic metabolic perturbations, and reduces tumor growth in three murine cancer models. The antitumor effects of chronic tirzepatide treatment in obese mice, on colon and breast cancer models are consistent with our findings in a murine model of endometrial cancer^30^, as well as a report by others on liver cancer^36^, and are further supported by reports that significant weight loss (at levels similar to those achieved with tirzepatide) in response to chronic or intermittent calorie restriction or bariatric surgery have anticancer effects in multiple obesity-related cancer models^44–46^. Moreover, emerging pharmacoepidemiologic evidence suggests that use of GLP1RA for the management of type 2 diabetes may lead to reduced risk of most obesity-related cancers^22^. Together, these findings suggest that use of tirzepatide is a promising interception strategy for obesity-associated cancers.

A novel finding in the present study is that tirzepatide exerts its anticancer effects by reversing CD8+ T cell-associated immunosuppression in the tumor microenvironment. We demonstrate that tirzepatide restored expression of transcriptomic signatures of antitumor immunity which were suppressed by obesity in multiple tumor models. In particular, a role for CD8+ TIL function on limiting MC38 tumor growth in tirzepatide-treated obese mice is established by the loss of tirzepatide’s beneficial effects upon systemic CD8+ T cell depletion with a neutralizing antibody. Obesity engenders an environment whereby CD8+ TILs are ineffective and hyporesponsive, resulting in rapid acceleration of tumor growth^6–8^. Indeed, obesity-driven defects in CD8+ TIL activation curtail the requirement for T cell extrinsic immune evasion to the point where tumors transplanted from obese mice into immunocompromised mice grow more slowly than those transplanted from never-obese control mice^8^. We demonstrate that tirzepatide protects against CD8+ TIL inactivity by reversing phenotypic markers of obesity-associated immunosuppression (decreased CD62L, TCF1; increased CD39, PD-1 relative to tumors from vehicle-treated DIO mice).

Direct effects of tirzepatide on MC38 colonic adenocarcinoma cells via agonism of incretin receptors to promote cytotoxicity are unlikely given the lack of expression of both GLP1R and GIPR on these cells, and the lack of response of these cells to tirzepatide across a wide dose range in vitro. However, direct effects of GLP1R agonism on CD8+TILs are plausible, given reports of GLP1R upregulation in activated TILs^39–41^ and our observation from single-cell sequencing data that a subset of activated CD8+ TILs express *GLP1R*. Further analysis of single-cell sequencing data indicated that *GLP1R*+ CD8+ TILs, relative to *GLP1R*-TILs, displayed uniform suppression of pathways involved in T cell activation and differentiation, chemotaxis, and interferon-mediated signaling, suggesting that direct effects of tirzepatide on CD8+ TILs are unlikely to explain its anticancer benefits. We further showed that agonism of GLP1R with tirzepatide in ex vivo splenocytes isolated from tumor-bearing DIO mice modestly suppressed CD8+ T cell effector function. Others have also indicated that GLP1R agonism on CD8+ T cells is immunosuppressive, as GLP1R on activated T cells acts as a negative costimulatory molecule akin to PD-1^39^. Congruently, GLP1R agonists exendin-4 and liraglutide suppress CD8+ T cell proliferation and cytokine production (consistent with our in vitro findings) to curb inflammation and ameliorate systemic and localized inflammatory conditions^39–41,47,48^. We thus conclude that direct engagement of the intended targets of tirzepatide, the GLP-1 and GIP receptors, on CD8+ TILs is not substantially involved in the antitumor effects of tirzepatide in obese mice.

There is a dearth of actionable targets for restoring CD8+ TIL functionality in the context of obesity^6–8,43^, and our finding that tirzepatide is able to do so, possibly via systemic metabolic reprogramming, may inform future studies. For example, circulating leptin, which is increased >9-fold in response to DIO but is largely normalized by tirzepatide treatment, is directly proportional to fat mass, and frequently implicated in suppressing antitumor immunity^9,42^. Leptin agonism of the leptin receptor (LEPR) on CD8+ T cells suppresses CD8+ TIL effector function, while CD8+ T cells lacking LEPR limit tumor growth in Rag2^-/-^ mice to a greater extent than LEPR-expressing T cells^9,42^. In addition, obesity-associated increases in leptin and other metabolic and inflammatory signals impairs antigen presentation by tumor-associated macrophages, thus preventing the proper activation and clonal expansion of antigen-reactive CD8+ TILs^43^.

We found that tirzepatide was ineffective at suppressing tumor growth in lean mice, suggesting resolution of underlying metabolic dysfunction (such as hyperleptinemia) is a predominant means through which tirzepatide protects against obesity-driven tumor growth. Consistent with this finding, Piening et al. recently demonstrated that weight loss through dietary restriction enhanced antitumor immunity in a B16-F0 mouse melanoma model, but a suboptimal semaglutide dosage that failed to resolve obesity-driven metabolic dysfunction did not^8^.

We conclude that the health benefits of incretin-mimetic drugs such as tirzepatide in individuals with obesity may include intercepting obesity-related cancers via improved antitumor immunity. While we show that tirzepatide clearly reduces the growth rates of established cancers in allograft models, additional laboratory studies are needed to further elucidate mechanism and to clarify if tirzepatide influences early steps of carcinogenesis. Long-term randomized clinical trials of incretin-mimetic drugs with cancer incidence or progression endpoints would be logistically challenging but clinical intervention studies with tissue and circulating biomarker endpoints are justified.

## Methods

### Mice

All studies were approved by the Institutional Animal Care and Use Committee at the University of North Carolina–Chapel Hill or the University of Michigan. C57BL/6J diet-induced obese (DIO; stock #380050) and age-matched control male mice (stock #000664) were purchased from the Jackson Laboratory at 10-16 weeks of age (Bar Harbor, ME). Female C57BL/6NCrl mice were purchased at 10 weeks of age (Charles River, Wilmington, MA). Mice were housed in a specific-pathogen-free facility with a 12 h light (7am-7pm)/12 h dark (7pm-7am) cycle, and cages were changed weekly to refresh diet and ALPHA-dri^TM^ bedding (Shephard Specialty Papers, Watertown, TN). Obese mice and age-matched control mice used for E0771 and MC38 tumor models were fed ad libitum a 60 kcal% fat diet (DIO; D12492 Research Diets, Inc, New Brunswick, NJ) or 10 kcal% fat diet (control; D12450J Research Diets, Inc.), respectively. CCR was implemented by providing mice a daily aliquot of a high protein diet (D15032801, Research Diets Inc.), to achieve a 30% reduction in caloric intake with adequate micronutrient consumption relative to controls. Mice used for Py230 tumor model underwent a modified diet-induced obesity program relying on a lower fat diet but housed in thermoneutral conditions to reduce energy expenditure^49^. These mice were housed at 28°C–31°C to limit adaptive thermogenesis and provided a 40 kcal% fat 39 kcal % sugar diet ad libitum (D15031601; Research Diets, Inc.). Body composition was assessed by magnetic resonance imaging (EchoMRI, Houston, TX). Mice were euthanized with CO_2_ inhalation followed by cervical dislocation or exsanguination while under isoflurane anesthesia.

### Tirzepatide dosage

After a minimum of 12 weeks on diet, DIO mice were randomized by body weight to remain obese or lose weight via tirzepatide (LY3298176, Selleck Chemicals, Houston, TX) while continuing the 60 kcal% fat diet. Mice were subcutaneously injected with either vehicle (5% DMSO/95% 40 mM Tris pH 8.0) or tirzepatide (dose ranging from 1-30 nmol/kg in pilot dose-finding study, 3-10 nmol/kg in tumor studies in male mice, 3-30 nmol/kg in female mice, and 3-75 nmol/kg in female mice housed at thermoneutrality; frequency ranging from daily to q.o.d.) for the remainder of each study. Injections occurred between 3:30-6:00 pm to proceed the onset of the dark cycle.

### Fasted blood glucose measurements

Mice were fasted for 5 h. Blood was collected from the tail vein, and glucose concentration was measured using a Contour Next EZ blood glucometer (Bayer, Jamestown, NC). For intraperitoneal glucose tolerance tests, mice were injected with a bolus of 1.5 mg/kg D-glucose, and blood was collected and read on a glucometer at 0, 15, 30, 60, 90, and 120 min.

### Circulating hormone and cytokine concentrations

Mice were fasted 4-5 h prior to collection of submandibular (prior to tumor injection) or cardiac blood (at study termination). Blood samples clotted at RT for 30-60 min and were centrifuged at 1000 x g for 15 min at 4 °C. Serum was aliquoted, flash frozen in liquid N_2_, and stored at -80 °C. Serum hormone concentrations were quantified using the Bio-Plex Pro Mouse Diabetes 8-plex and Adiponectin assays (Bio-Rad Laboratories, Hercules, CA), and IGF-1 mouse Luminex discovery assay (R&D Systems, Minneapolis, MN) using the Luminex platform. In the study using only lean mice, a smaller marker panel was used that included mouse insulin (Crystal Chem, Elk Grove Village, IL), mouse leptin (Crystal Chem), and total mouse IGF-1 (Ansh Labs, Webster, TX), each measured by ELISA according to manufacturers’ instructions.

### Tumor cell transplantation

MC38 (Kerafast, Boston, MA) and E0771 (CRL-3461, ATCC, Manassas, VA) cells were maintained at 37 °C with 5% CO_2_ in Dulbeco’s Modified Eagle Medium containing 4.5 g/L glucose (Gibco, Carlsbad, CA) and supplemented with 10 mM HEPES buffer (Gibco), 10% fetal bovine serum (VWR International, Radnor, PA), 2 mM glutamine (Gibco), and 1% penicillin streptomycin (Gibco). Py230 cells (provided by Nicholas Webster and Lesley Ellies, University of California San Diego) were maintained in F12K Nutrient Mixture (Corning 10-025-CV) supplemented with 5% Fetal Clone II (Cytiva SH30066.03), 50 µg/mL gentamycin (Sigma-Aldrich G1397), 2.5 µg/mL amphotericin B (Sigma-Aldrich A2942), and 1 µL/mL MITO+ Serum Extender (Corning 355006) at 37 °C and 5% CO_2_. Cells were routinely tested for the presence of mycoplasma using the Universal Mycoplasma Detection Kit (ATCC). Mice were subcutaneously injected with 2.5 x 10^5^ MC38 cells into the right flank. Mammary cancer tumors were achieved by orthotopically injecting 3.5 x 10^4^ E0771 cells or 1 x 10^4^ Py230 cells in 2 mg/mL Matrigel (Corning 356234) into the 4^th^ mammary fat pad. Tumor growth was monitored every 2-3 days using digital calipers. Tumor volume was calculated as D x d^2^ x 0.5 where D is the diameter of the longer dimension and d is the diameter perpendicular to D^50^.

### CD8+ cell depletion

Mice were intraperitoneally injected with either 200 μg rat IgG2b isotype control (#BE0090, clone LTF-2; Bio X Cell, Lebanon, NH) or 200 μg αCD8α (#BE0061, clone 2.43; Bio X Cell). Relative to the day of MC38 tumor cell injections, isotype control and αCD8α antibodies were injected on days -2, 0, 2, and every 4 days thereafter.

### High-dimensional flow cytometry

Tumors were dissociated using the Miltenyi Biotec (Bergisch Gladbach, Germany) mouse tumor dissociation kit and gentleMACs dissociator according to manufacturer’s instructions but reducing enzyme R to 20% of suggested volume. Leukocytes were enriched using Percoll density gradient centrifugation. Spleens were mechanically digested and filtered through a 70 μm cell strainer, and red blood cells were lysed with ammonium-chloride-potassium buffer (Gibco).

For T cell phenotyping, cells were stained in PBS for 30 min at 4 °C with Zombie NIR viability dye (1:7000; Biolegend, San Diego, CA), CD45-BV510 (1:1000; BD Biosciences, Franklin Lakes, NJ), CD8⍺-Spark NIR 685 (1:1000; Biolegend), CD4-APC Fire 810 (1:1000; Biolegend), CD11b-Alexa Fluor 532 (1:5000; Invitrogen), TCRβ-BV570 (1:100; Biolegend), CD44-Brilliant Violet 785 (1:400; Biolegend), CD62L-BV421 (1:400; BD Biosciences), PD-1-BB700 (1:200; BD Biosciences), TIM-3-BV711 (1:200; Biolegend), LAG-3-APC-eFluor 780 (1:200; Invitrogen), SlamF6-APC (1:200; Invitrogen), CD39-Super Bright 436 (1:200; Invitrogen), CX3CR1-PE-Fire 640 (1:200; Biolegend), CD69-PE-Cy5 (1:600; Biolegend), GITR-BV650 (1:400; BD Biosciences), CD27-BV750 (1:200; BD Biosciences), and CD16/CD32-purified FcR block (Invitrogen). Cells were washed and fixed overnight using the Foxp3/Transcription Factor Fixation/Permeabilization Concentrate and Diluent kit (Invitrogen). Cells were then washed and stained with T-bet-eFluor 450 (1:100; Invitrogen), CTLA-4-PE-Dazzle 594 (1:400; Biolegend), Ki67-BV605 (1:400; Biolegend), TOX-PE (1:100; Miltenyi Biotec), TCF-1-PE-Cy7 (1:100; Cell Signaling Technology, Danvers, MA), Granzyme B-Alexa Fluor 700 (1:200; Biolegend), and BCL-2-Alexa Fluor 647 (1:400; Biolegend) for 3 h at RT.

For splenic and intratumoral CD8+ T cell depletion assessment, cells were incubated with TruStain FcX (Biolegend) in PBS for 15 min at 4 °C. Cells were stained with LIVE/DEAD fixable blue dead cell stain (1:1600; Invitrogen), CD45-BUV395 (1:300; BD Biosciences), CD4-BUV805 (1:200; BD Biosciences), CD8⍺-Alexa Fluor 594 (1:200; Biolegend), and CD3-Alexa Fluor 700 (1:200; Biolegend). To assess CD8+ T cell enrichment from splenocyte preparations, cells were additionally stained with rat anti-mouse CD11c-BUV737 (1:100; BD Biosciences), CD11b-PerCP-eFluor 710 (1:100; Invitrogen), and CD19-PerCP (1:75; Biolegend). Cells were fixed using a cytofix/cytoperm fixation/permeabilization kit (BD Biosciences).

Events were collected on a Cytek Biosciences (Fremont, CA) Aurora or Northern Lights full spectrum flow cytometer. Data were analyzed using FlowJo (v.10.8.1; Ashland, OR) and OMIQ software (Dotmatics, Boston, MA).

### CD8+ splenocyte in vitro activation and expansion

CD8+ T cells were isolated from spleens from tumor-bearing obese mice by negative magnetic selection (STEMCELL Technologies, Vancouver, Canada). Isolated cells were activated with 5 μg/ml plate-bound αCD3, 2 μg/ml soluble αCD28 (eBioscience, San Diego, CA), and 200 units/ml IL-2 (TECIN) and treated with 0-1000 nM tirzepatide for 3 days in RPMI supplemented with 10% FBS, 300 mg/L L-glutamine, 100 units/mL penicillin, 100 μg/mL streptomycin, 1mM sodium pyruvate, 100 μM NEAA, 1 mM HEPES, 55 μM β-mercaptoethanol, and 0.2% plasmocin mycoplasma prophylactic. On day 3, media was replaced as at day 0, except lacking αCD3 and αCD28. Cytokine production was assessed on day 5 following 5 hours of restimulation with PMA/ionomycin [Cell Stimulation Cocktail (Thermo Fisher Scientific) at 1:500].

### Cell viability assays

5 x 10^2^ MC38 or E0771 cells were seeded in complete media in a 96-well plate. Media was replaced 4 h after seeding with complete media containing 0-500 nM tirzepatide. Media (with or without tirzepatide) was replaced once after 44 h. 72 h after seeding, cells were incubated in 3-(4,5-dimethylthiazol-2-yl)-2,5-diphenyltetrazolium bromide (MTT) reagent for 75 min. Formazan was solubilized in 100 μl DMSO, and the relative absorbance at 570 nm subtracted from background absorbance at 690 nm was measured on a Cytation3 plate reader (BioTek Instruments; Winooski, VT) to assess cell viability.

### Immunohistochemistry

Tissues were fixed in 10% neutral-buffered formalin for 24 h and stored in 70% ethanol at RT. Paraffin-embedded tissue was baked overnight, deparaffinized, rehydrated, and underwent heat-induced epitope retrieval using Borg Decloaker (Biocare Medical, Pacheco, CA) and a decloaking chamber (120 °C for 30 sec followed by 90 °C for 10 sec). Slides were incubated in 3% hydrogen peroxide for 10 min at RT and then permeabilized and blocked in 5% normal goat serum for 1 h at RT. Slides were incubated overnight at 4 °C with ɑ-GLPR (1:500, rabbit, ab218532; Abcam, Cambridge, United Kingdom). Slides were then sequentially incubated with biotinylated ɑ-rabbit IgG (1:500, goat, #111-065-144; Jackson ImmunoResearch Labs, West Grove, PA) for 1 h at RT and an ABC Elite reaction kit (1:50, #PK-6100; Vector Laboratories, Burlingame, CA). Positive signal was detected with diaminobenzidine (Thermo Fisher Scientific), and slides were counterstained with hematoxylin.

### Cancer Cell Line Encyclopedia gene expression analysis

Human cancer cell line gene expression data was accessed from the Broad Institute’s Cancer Cell Line Encyclopedia through the Depmap portal^36^. “Batch corrected Expression Public 24Q2” was downloaded with metadata. A cancer cell line *GLP1R* or *GIPR* expression of log2(TPM+1) >2 was deemed to be appreciable^37^.

### Bulk RNA sequencing

RNA was isolated from whole tumors using the RNeasy micro kit (Qiagen), quantified using a Qubit 2.0 fluorometer (Life Technologies, Carlsbad, CA), and assessed for integrity using a TapeStation 4200 (Agilent Technologies, Santa Clara, CA). NEBNext Ultra II RNA libraries were prepared (New England Biolabs, Ipswitch, MA) with an ERCC RNA spike-in (Thermo Fisher Scientific) and sequenced using an Illumina HiSeq with a 2 x 150 bp paired-end configuration targeting >25 million reads per sample. Adapters and low-quality reads were removed with Trimmomatic^51^. Spliced Transcripts Alignment to a Reference (STAR) was used to align reads to the GRCm38 mouse genome^52^. Genes with fewer than 10 read counts in 5 or more samples were removed from all downstream analyses^53^.

### Single-cell RNA sequencing analysis

Publicly available scRNA-seq data of TIL isolated from control and DIO mice bearing B16-F0 tumors was accessed at GSE245657^8^ and processed in R (v4.2.2) using Seurat (v5.0.3)^54^. Cells were filtered to include cells with 500-6,500 total reads and with mitochondrial gene expression <10%. 2,491 cells from control mice and 2,373 cells from DIO mice were analyzed. UMAP embeddings were generated using the first 15 principal component analysis components with resolution of 1.3. Canonical correlation analysis was used to integrate control and DIO UMAP embeddings. Redundant clusters were collapsed by Euclidean distance of mean-cluster-wise gene expression using cutree (v4.3.2). Cluster identity was manually assigned from curated gene lists^55^. Any cell with non-zero expression of *GLP1R* was considered *GLP1R* expressing (*GLP1R*+) (∼1.5% overall).

### Statistical analysis

Statistical analyses were performed using GraphPad Prism (v10.2.2, GraphPad Software Inc., La Jolla, CA) and R (v.4.2.2). Unpaired t-tests compared two groups. One-way ANOVAs with Tukey’s post hoc test compared more than two groups. Repeated measures two-way ANOVAs with Šídák’s multiple comparisons tests compared two groups across time. Repeated measures one-way ANOVA with Dunnett’s multiple comparison test assessed percent viability of tirzepatide-treated cells relative to vehicle-treated cells for the MTT assay. Correlation between tumor burden and body weight was determined by Spearman correlation. Tumor growth over time was assessed using linear regression with false discovery rate (FDR) q-value correction.

Differential abundance analysis for high-dimensional flow cytometry analysis was performed using edgeR. Ranked gene expression of *GLP1R*+ cells relative to all other cells from clusters containing at least 1 *GLP1R*+ cell was subject to gene set enrichment analysis^56^ using GOBP gene sets^57^. More than 10^7^ permutations were performed to ensure stable p values, which were subject to correction using the Benjamini-Hochberg procedure. GSEA of bulk transcriptomic profiling was performed using Hallmark gene sets and 10^3^ permutations. Outliers for hormone concentrations and MFIs for phenotypic markers were identified using the ROUT method (Q=5%) and removed. p<0.05 was considered statistically significant.

## Acknowledgements

The authors acknowledge the Preclinical Research Unit (supported in part by an NCI Center Core Support Grant CA16086 to the Lineberger Comprehensive Cancer Center), the Histology Research Core Facility, and the Genomics and Energy Metabolic Core (supported in part by NIDDK under award number P30DK056350) for their technical assistance in completing this study. This manuscript represents an update of bioRxiv 2023.06.22.546093 (doi: 10.1101/2023.06.22.546093) and bioRxiv 2024.01.20.576484 (doi: 10.1101/2024.01.20.576484).

## Author contributions

*Conceptualization*: E.M.G., M.F.C., G.T.C., B.P.R., V.A.K., E.D.G., J.E.T., M.P., S.D.H.; *Methodology*: E.M.G., M.F.C., G.T.C., B.P.R., V.A.K., E.M.P., E.D.G., J.E.T., M.P., S.D.H.; *Investigation*: E.M.G., M.F.C., G.T.C., B.P.R., A.N.H., T.L.M., V.A.K., A.L.K., E.G.B., K.E.S., J.Z., Y.W., H.M.M., C.E.G., F.C., R.X., C.A.H.; *Formal analysis*: E.M.G., M.F.C., G.T.C., B.P.R., J.E.T., S.D.H.; *Visualization*: E.M.G., M.F.C., G.T.C., B.P.R.; *Writing—original draft:* E.M.G., G.T.C., J.E.T., S.D.H.; *Writing—review & editing:* E.M.G., M.F.C., G.T.C., B.P.R., A.N.H., T.L.M., V.A.K., A.L.K., E.G.B., K.E.S., J.Z., Y.W., H.M.M., C.E.G., F.C., R.X., C.A.H., E.M.P., J.R., E.D.G., J.E.T., M.P., S.D.H.; *Supervision*: E.D.G., J.E.T., M.P., S.D.H.; *Funding acquisition*: E.D.G., J.E.T., M.P., S.D.H.

