## Supplementary data for "Tirzepatide restricts obesity-related tumor growth by reversing metabolic dysregulation and rescuing CD8+ T cell function"

**Supplementary Figure 1. Tirzepatide (TZP) induces weight loss in obese mice in a dose-dependent manner.** **(A)** Percent weight change over 4 weeks (n=6/6/6/6) and **(B)** after 4 weeks of daily TZP treatment at indicated concentrations (1-10 nmol/kg) (n=6/6/6/6). **(C)** Cumulative food intake (n=4/4/4/4) and **(D)** fasting blood glucose concentration (n=5/5/5/6) of mice treated daily at indicated dose. **(E)** Percent weight change over 4 weeks (n=6/6/6/6) and **(F)** after 4 weeks of every other day (q.o.d.) of TZP treatment at indicated concentrations (3-30 nmol/kg) (n=6/6/6/6). **(G)** Cumulative food intake (n=4/4/4/4) and **(H)** fasting blood glucose concentration of mice treated q.o.d. at indicated TZP dose (n=5/6/6/6). **(I)** Percent weight change over 4 weeks (n=6/6/6/6) and **(J)** after 4 weeks of TZP treatment thrice weekly (Monday/Wednesday/Friday; MWF) at indicated concentrations (3-30 nmol/kg) (n=6/6/6/6). **(K)** Cumulative food intake (n=4/4/4/4) and **(L)** fasting blood glucose concentration (n=5/6/6/6) of mice treated MWF at indicated dose. The same group of 6 mice receiving the DIO diet and daily vehicle treatment served as the reference group for each TZP dosing strategy.

Data are represented as mean ± SEM. Statistical significance determined by repeated measures two-way ANOVA with Tukey's multiple comparisons test (A, E, I), one-way ANOVA with Tukey's multiple comparisons test (B, D, F, H, J, L), or area under the curve (AUC) (C, G, K).

**Supplementary Figure 2. Tirzepatide (TZP) limits tumor growth in breast cancer models.** **(A)** Body weight in E0771 bearing control, DIO, TZP, and CCR mice (n=14/15/13/15). **(B)** Fat mass and (**C**) % fat mass in E0771 bearing DIO and TZP mice (n=10/10). (**D**) E0771 tumor weight in control, DIO, TZP, and CCR mice (n=13/15/13/13). **(E)** Py230 tumor volume in DIO and TZP mice (n=6/9). **(F-G)** Spearman correlation between E0771 (n=57) and Py230 (n=15) tumor burden and body weight. **(H)** Immune-related gene sets across tumors from DIO and control or TZP mice in MC38 (n=6/6/4), E0771 (n=4/6/6), and Py230 (n=6/10) models.

Data are represented as mean ± SEM. Statistical significance determined by one-way ANOVA with Tukey's multiple comparisons test (A,D), unpaired two-sided Student's t-test (B-C,E), or Spearman correlation (F-G).

**Supplementary Figure 3. Diet drives splenic T cell phenotype.** **(A)** CD62L (n=8/8/8), CD44 (n=8/7/8), Ki67 (n=8/8/8), BCL-2 (n=8/8/8), CD27 (n=8/8/8), BODIPY (n=8/8/8), CX3CR1 (n=8/7/8), TOX (n=8/8/8), TIM-3 (n=8/8/8), PD-1 (n=8/8/8), and TCF-1 (n=7/8/8) MFIs. **(B)** % CD8+ splenic T cells of live CD45+/CD3+ cells 14 days post MC38 subcutaneous injection (n=3/3). **(C)** % CD8+ tumoral T cells of live CD45+/CD3+ cells 14 days post MC38 subcutaneous injection (n=3/3). **(D)** % CD8+ tumoral T cells of live CD45+/CD3+ cells at termination of study (n=24/24).

Data are represented as mean ± SEM. Statistical significance determined by one-way ANOVA with Tukey's multiple comparisons test (A) or unpaired two-sided Student's t-test (B-D).

**Supplementary Figure 4**. ***Glp1r*+ CD8+ T cells exhibit reduced features of activation**. UMAPs visualizing unsupervised clustering of published scRNA-seq data (GSE245657) of **(A)** CD8+ TILs isolated from B16-F0 tumors and **(B)** Glp1r+ cells. Ice blue dots indicate *Glp1r*+

cells isolated from control mice; green dots indicate *Glp1r+* cells isolated from obese mice. Numbers indicate the number of *Glpr+* cells relative to total cells sequenced from each diet group. **(C)** Heatmap showing relative expression of genes used to define each scRNA-seq cluster. **(D)** Gene set enrichment analysis using GOBP gene sets in activated *Glpr1-* vs *Glp1r+* cells from scRNA-seq data. **(E)** Representative histograms and quantification of in vitro **(F)** IFNγ production (n=3/3/3) and **(G)** TNFα production (n=3/3/3) in splenic T cells isolated from tumor-bearing obese mice and subsequently activated and treated with TZP in vitro at indicated doses for 5 days. Dots of the same color indicate same biological replicate.

Data are represented as mean ± SEM. Statistical significance determined by repeated measures one-way ANOVA with Dunnett's multiple comparisons test (F-G).

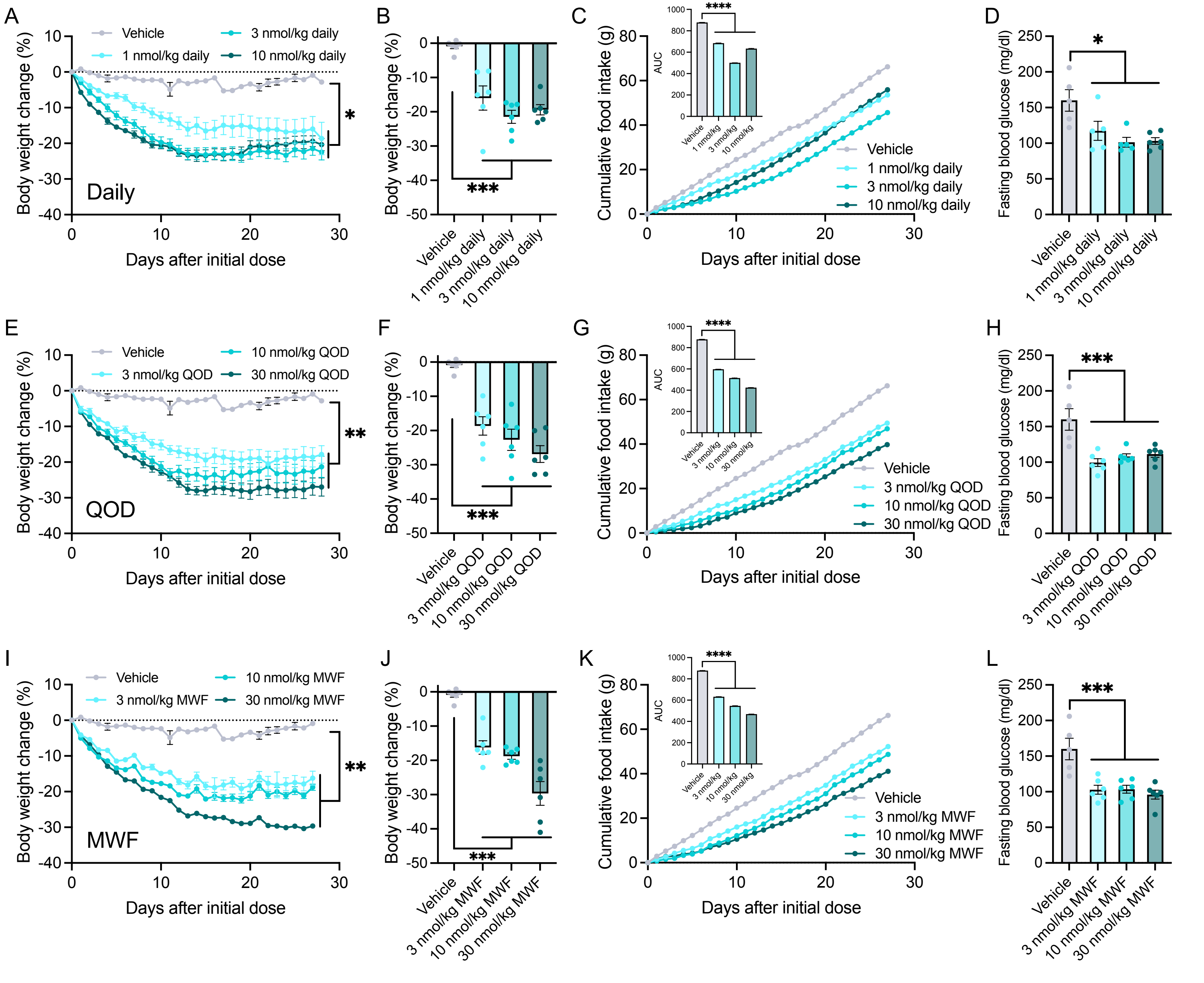


Supplemental Figure 1

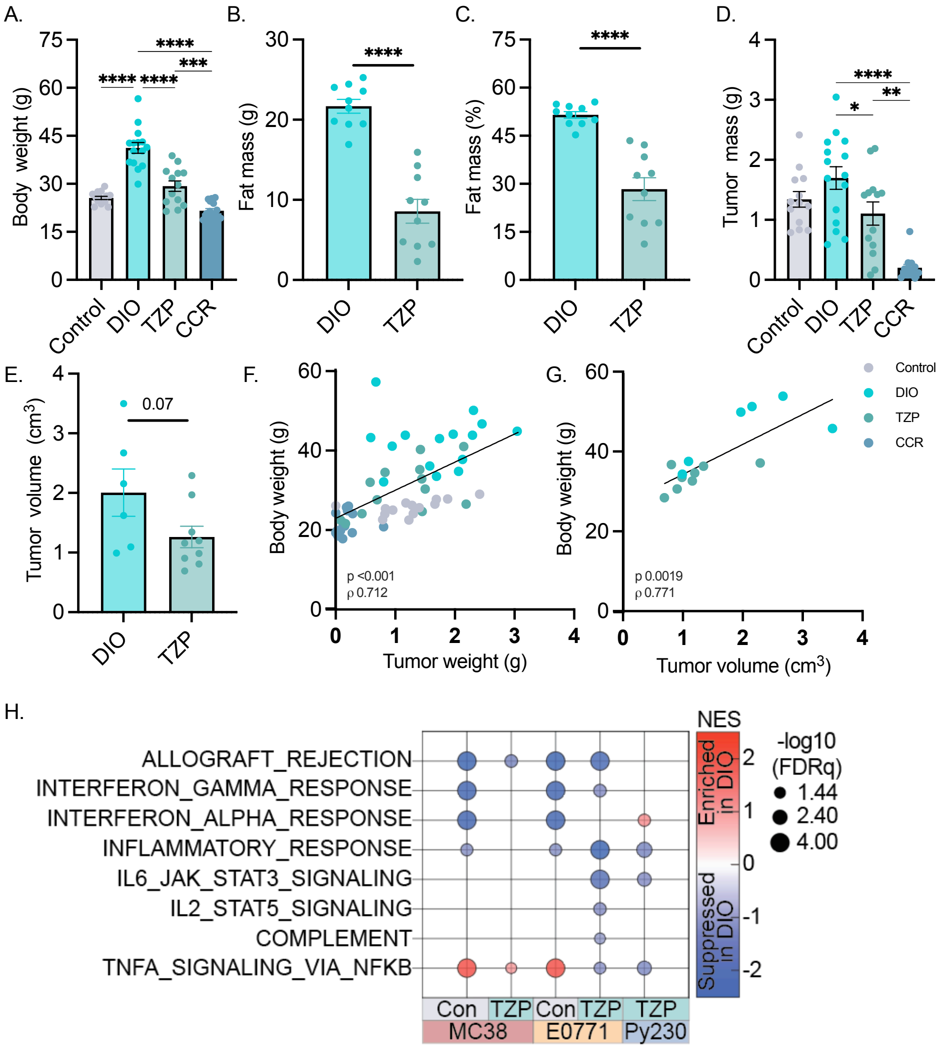


Supplemental Figure 2

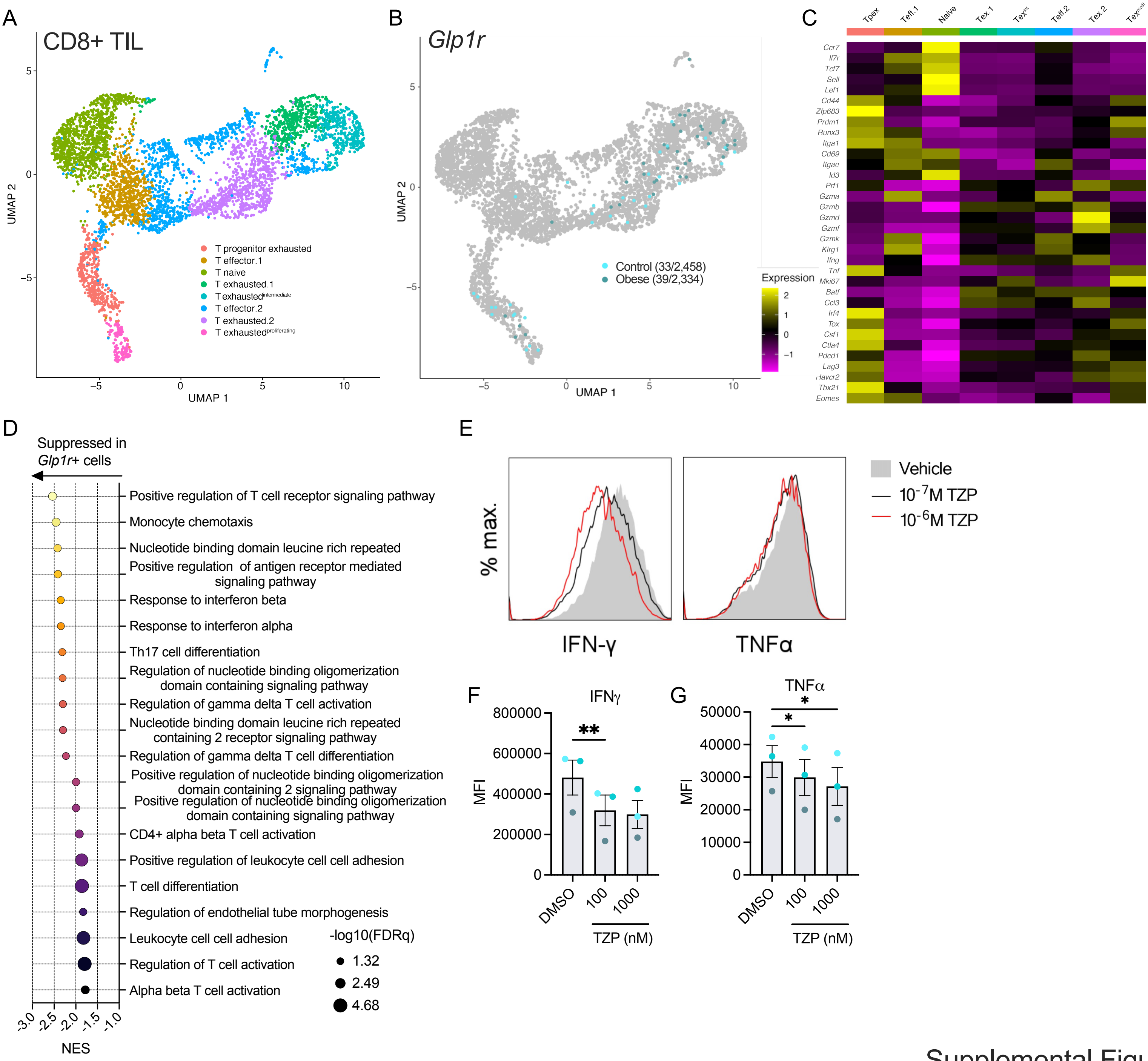


Supplemental Figure 3

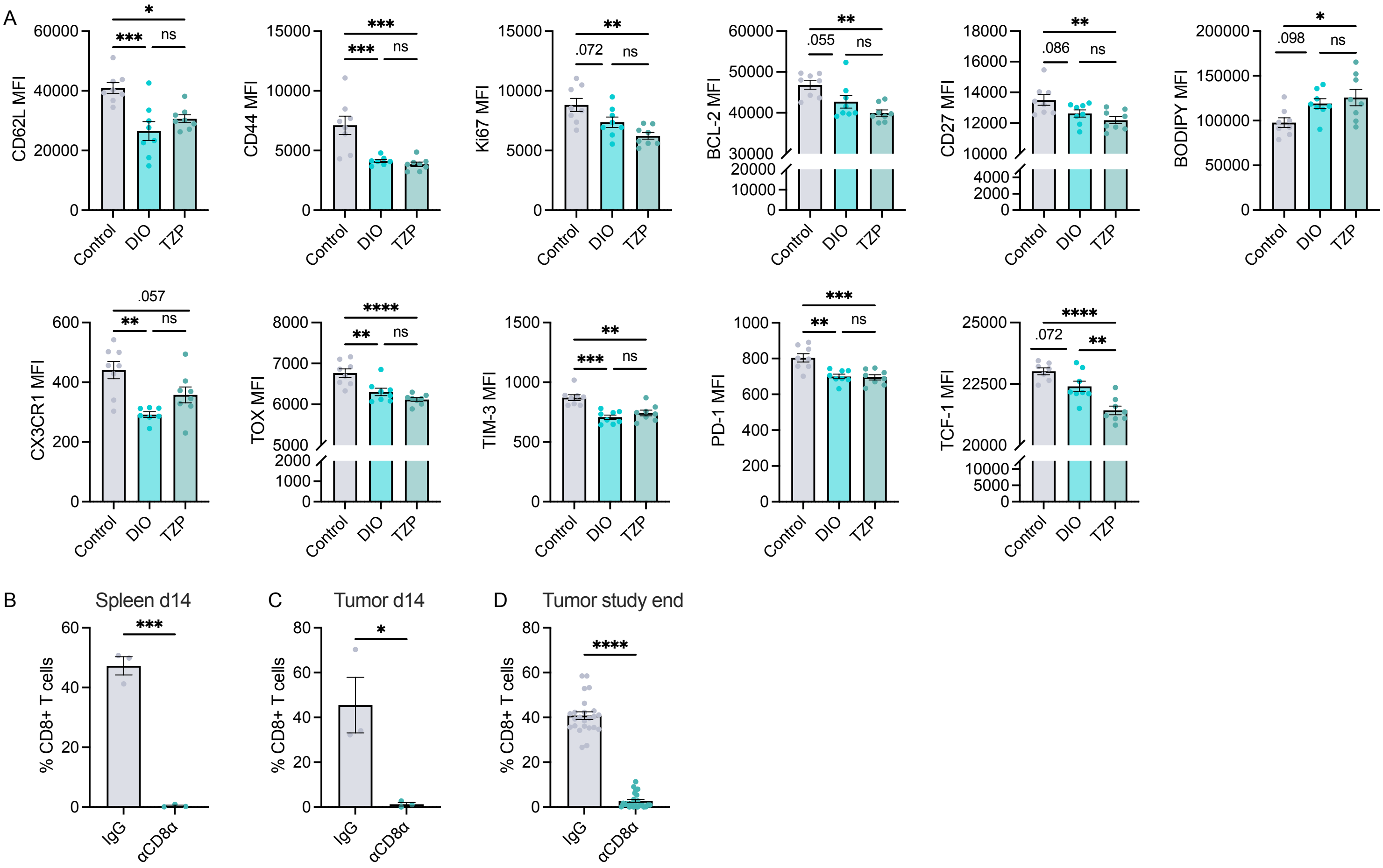


Supplemental Figure 4

**Table S1.** Fasting concentrations of circulating glucose and selected metabolic hormones.

| Analyte | Control | DIO | TZP | CCR |
|---|---|---|---|---|
| Adiponectin (μg/ml) | 20.0 (7.5)† | 23.1 (6.8)† | 28.7 (13.4)† | 95.0 (40.0)*# |
| Ghrelin (ng/ml) | 15.5 (6.5)† | 10.7 (3.0)† | 16.2 (5.1)† | 27.4 (12.1)*# |
| GIP (ng/ml) | 1.9 (0.4) | 1.8 (0.6) | 2.0 (0.5) | 2.0 (0.3) |
| GLP-1 (pg/ml) | 124.2 (66.2) | 108.6 (48.0) | 100.8 (48.2) | 130.2 (36.7) |
| Glucose (mg/dl) | 127.6 (15.0)† | 149.5 (18.5)*#† | 124.5 (15.5) | 107.6 (18.8)* |
| Glucagon (pg/ml) | 847.6 (279.7) | 702.0 (185.3)† | 696.3 (94.0)† | 972.2 (252.7)# |
| IGF-1 (ng/ml) | 30.5 (5.3) | 53.8 (18.2)*#† | 40.0 (10.3) | 36.5 (7.9) |
| Insulin (ng/ml) | 2.6 (0.9) | 4.5 (1.2)*#† | 2.6 (0.5) | 3.1 (1.6) |
| Leptin (ng/ml) | 0.6 (0.2)# | 11.9 (4.4)*#† | 3.8 (2.5)* | 1.6 (1.2) |
| PAI-1 (ng/ml) | 2.6 (1.4)#† | 1.7 (0.6) | 1.1 (0.5)* | 1.0 (0.4)* |
| Resistin (ng/ml) | 89.8 (25.2) | 65.5 (15.3)*† | 69.0 (17.7) | 90.0 (16.5) |

Mean (SD). Blood glucose measured on week 9 of weight loss intervention. Serum hormones assayed following 13-week intervention. One-way ANOVA with Tukey's post hoc test. *p≤0.05 vs. Control; #p≤0.05 vs. TZP; †p≤0.05 vs. CCR.

CCR, chronic calorie restriction; DIO, diet-induced obesity; GIP, glucose-dependent insulinotropic polypeptide; GLP-1, glucagon-like peptide 1; IGF-1, insulin-like growth factor 1; PAI-1, plasminogen activator inhibitor 1; TZP, tirzepatide.
